# Age-Dependent Effects of Macrophage Derived Cathelicidin Antimicrobial Peptide on Bone Fracture Repair

**DOI:** 10.64898/2026.09.29.755446

**Authors:** Tuyet Nguyen, Yasuhito Yahara, Eijiro Shimada, Vijitha Puviindran, Puviindran Nadesan, Koji Ishikawa, Choiselle Marius, Benjamin Alman

## Abstract

The pace of fracture repair declines with age, increasing the risk of delayed union and nonunion. Delayed union and nonunion are associated with significant morbidity and even mortality with few effective non-surgical interventions. Here, we identify cathelicidin antimicrobial peptide (CAMP) as an age-dependent regulator of bone repair. Single-cell RNA sequencing and immunohistochemistry demonstrated higher CAMP expression during fracture healing in young compared with aged mice. Global or myeloid lineage-depletion of *Camp* delayed fracture repair in young mice, whereas *Camp* depletion in aged mice increased callus bone formation. Heterochronic bone marrow transplantation using the conditional genetic knockout further implicated hematopoietic-lineage derived cells in these effects and demonstrated that myeloid-specific *Camp* expression is important in the rejuvenation of the pace of fracture healing. CAMP promotes osteogenic differentiation through P2X7 receptor (P2X7R)-dependent signaling, influencing the downstream regulation of NLRP3 expression. NLRP3 expression is also correlated with osteogenesis following *Camp* modulation in both young and old mice, supporting CAMP-P2X7R-NLRP3 signaling as a regulator of osteogenesis. Together, these findings identify CAMP as a hematopoietic-derived regulator of bone repair whose function is age-dependent, promoting fracture healing in young animals while constraining it in aged animals.

## Introduction

Fracture risk and complications increase with age, creating a growing health burden as the U.S. population over 65 expands^1, 2^. Older individuals are more likely to experience delayed union or non-union, complications associated with substantial morbidity and, in severe cases, increased mortality^3-6^. Bone fracture repair proceeds through four overlapping stages: inflammation, cartilaginous callus formation, hard callus formation, and bone remodeling^7^. With advanced age, fracture healing becomes characterized by delayed repair, prolonged cartilage persistence, and increased fibrosis^8-10^.

Aging also promotes myeloid-biased hematopoiesis^11, 12^. Changes in hematopoiesis can also be associated with reduced osteogenesis^13, 14^. Previous studies have demonstrated that myeloid cells, themselves, also undergo age-related functional changes^15, 16^. These aged myeloid cells are associated with impaired fracture healing, characterized by reduced bone volume and weaker mechanical properties^17^. Since myeloid cells are present during the inflammatory phase of fracture repair, its direct interaction with other cells or through secreted factors are observed to influence the repair process. However, how age-related changes in hematopoietic cells influence the fracture healing process remains incompletely understood.

Cathelicidin antimicrobial peptide (CAMP), also known as LL-37 in humans, is an evolutionarily conserved host-defense peptide produced primarily by innate immune cells, including neutrophils and macrophages. Beyond its direct antimicrobial activity, CAMP regulates inflammation, immune cell recruitment, angiogenesis, and tissue repair^18-20^. Its biological effects are context dependent and are mediated through several receptors and signaling pathways, including the purinergic receptor P2×7^19, 21-23^. CAMP also contributes to the resolution of diverse tissue injuries, from skin wound healing to thrombosis, likely through modulation of cytokine responses and promotion of immune cell migration^24-26^. Recombinant CAMP supplementation has been shown to promote osteogenesis in a bone defect model^20, 27^. However, the role of endogenous CAMP in fracture repair has not been rigorously evaluated *in vivo*, particularly in the context of aging.

## Results

### Cathelicidin antimicrobial peptide is expressed at higher levels following fracture in young mice compared with aged mice

Fracture healing in older animals is characterized by a slower pace of healing, compared to their younger counterparts. Aging is also associated with a myeloid differentiation bias of hematopoietic stem cells, resulting in increased production of common myeloid progenitors, granulocyte-monocyte progenitor, and megakaryocyte-erythroid progenitors ^8, 12, 28^. The impact of these age-related changes on bone repair remains poorly understood. Therefore, we sought to characterize age-associated transcriptomic changes in the context of bone injury. We performed single-cell RNA sequencing (scRNA-seq) analysis of femoral bicortical drill-hole injuries in young (2-month-old) and aged (20-month-old) mice. 3 days post injury, during the inflammation phase, hematopoietic cells were isolated and pooled according to age (Fig 1a).

**Figure 1.**
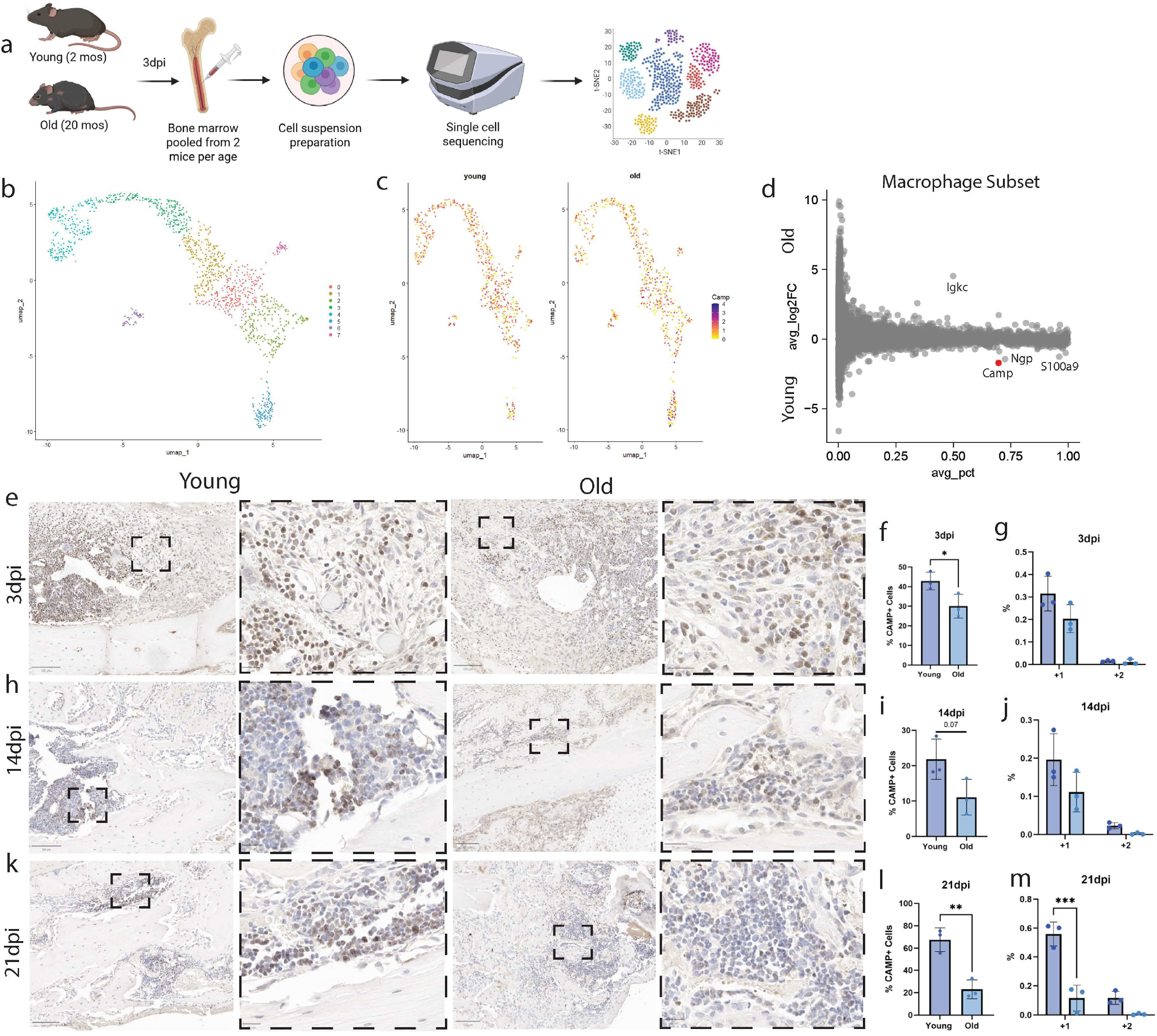
Cathelicidin antimicrobial peptide is expressed at higher levels following fracture in young mice compared with aged mice. a) Schematic of the preparation of the single cell data from young mice (2 months) and old mice (20 months) after a bicortical femoral drill hole injury. b) Integrated young and old macrophages at 3 days post-drill hole. c) Feature plot demonstrating Camp expression within the young vs old macrophage populations. d) MA plot of the macrophage population. e) Immunohistochemistry of CAMP in young and old mice 3 days post-tibial fracture. f) Quantification of CAMP+ cells 3dpi. g) Intensity of DAB staining of CAMP+ cells 3dpi. h) IHC of CAMP in young and old mice 14dpi. i) Quantification of CAMP+ cells 14dpi. j) Intensity of DAB staining of CAMP+ cells 14dpi. k) IHC of CAMP in young and old mice 21dpi. l) Quantification of CAMP+ cells 21dpi. m) Intensity of DAB staining of CAMP+ cells 21dpi. Results are presented as means ± SD. Statistical significance was determined using 2 tailed Welch’s t-test. n ≥ 3, *p < 0.05, **p < 0.01, ***p < 0.001

After quality control and filtering measures following the Seurat pipeline, young and old samples were then integrated together for downstream analysis (Suppl Fig 1a). Cell populations were annotated based on feature genes and cross-referenced with the CellMatch database^29^. The macrophage population was subsequently isolated to identify age-associated differences in gene expression (Fig 1b). Among the differentially expressed genes, *Camp* exhibited the most pronounced age-associated difference and was expressed in a greater proportion of young macrophages (Fig 1d). Immunohistochemistry (IHC) confirmed that the abundance of CAMP+ cells within the fracture callus was significantly greater in young than aged mice throughout the healing process, from 3-to 21-days post-fracture (Fig 1f, i, l). From these positive cells, we also observe that young mice express CAMP at a higher intensity than their older counterparts (Fig 1g, j, m).

### Global and myeloid-specific Camp depletion delays fracture healing

To determine *Camp*’s function within the context of bone healing, we used a global *Camp* depletion transgenic line. Using the same drill-hole injury model employed for the single-cell expression analysis, young (4-6 months) mice were collected 7 days after injury (Fig 2a). microCT analysis revealed a decrease in bone volume over total volume (BV/TV) and apparent density in the global *Camp* depletion mutants compared to their control siblings (Figure 2b-c). With the uninjured contralateral limb, we also examined the same parameters and observed no differences, confirming that the observed deficits in the microCT parameters are due to injury and not the genetic depletion (Supp Fig 2c-d). Then, we used a stabilized mid-diaphyseal tibial osteotomy (fracture) model to examine the role of *Camp* in endochondral bone repair under conditions more representative of a human fracture injury (Figure 2d-e). Following tibial fracture, *Camp*-depleted mice exhibited decreases in the BV/TV and the apparent density at both 14- and 21-days post injury (dpi), accompanied by an increase in the callus volume at 21dpi (Fig 2f-h). These results exemplify that the fracture callus in *Camp* deficient mice is composed of more callus tissue that has not fully ossified into bone, suggesting a delay in healing (Fig 2j-k).

**Figure 2.**
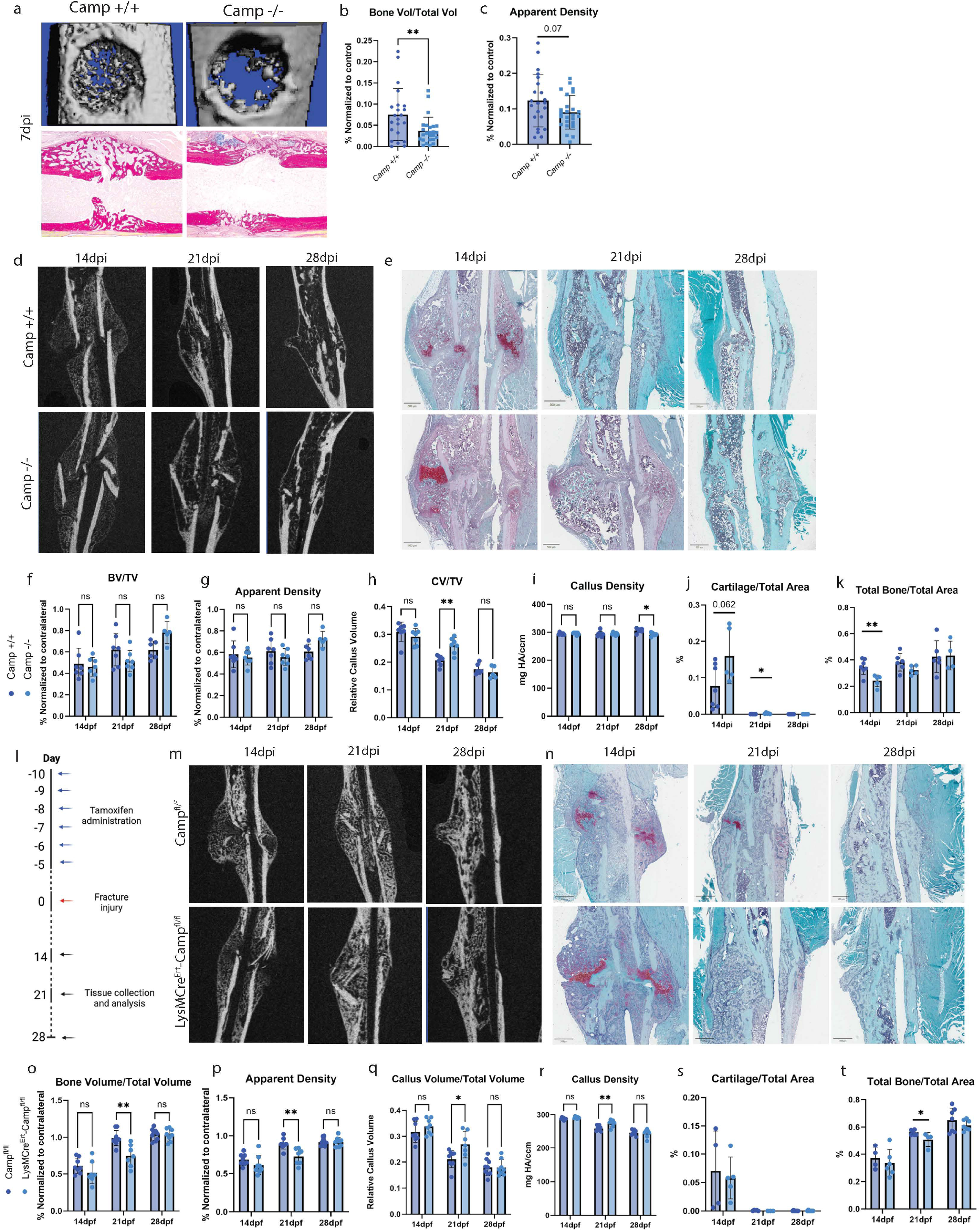
Global and myeloid-specific *Camp* depletion delays fracture healing. a) Representative images of young (4-6 months) *Camp* global knockouts 7 days after a femoral bicortical drill hole injury. b) relative bone volume (BV/TV) quantification 7dpi. c) Apparent density quantification 7dpi. d) Representative microCT images of young *Camp* global knockouts 14, 21, and 28 days after a stabilized tibial osteotomy. e) Representative Safranin-O/Fast Green images. f-i) Quantification of BV/TV, apparent density, relative callus volume (CV/TV), and callus density of young Camp global knockout mice. j-k) Quantification of cartilage and total bone area over total callus area. l) Schematic of tamoxifen administration and fracture collection timepoints for the *LysMCre*^*Ert*^*-Camp*^*fl/fl*^ mice. m) Representative microCT images of young myeloid-specific *Camp* knockouts 14, 21, and 28 days after a stabilized tibial osteotomy. n) Representative Safranin-O/Fast Green images. o-r) Quantification of BV/TV, apparent density, CV/TV, and callus density of young myeloid-specific *Camp* knockout mice. s-t) Quantification of cartilage and total bone area over total callus area. Results are presented as means ± SD. Statistical significance was determined using 2 tailed student’s t-test. n ≥ 3, *p < 0.05, **p < 0.01

Because macrophages exhibited the largest age-associated change in cellular abundance in our scRNA-seq dataset, we then determined whether myeloid-derived *Camp* contributes to the fracture healing phenotype. We conditionally depleted *Camp* in myeloid cells using the *LysMCre*^*Ert*^*-Camp*^*fl/fl*^ model. Tamoxifen was administered at 75mg/kg through intraperitoneal injections once every 24 hours for six consecutive days. The mice were then allowed to recover for five days before undergoing the same tibial fracture procedure as the global *Camp* depleted mice (Fig 2l). With this tamoxifen schedule, we demonstrate approximately 70% lower *Camp* mRNA levels from CD45+F4/80+ cells collected using fluorescence activated cell sorting at the time of fracture (Suppl Fig 2j).

Following fracture, mice were studied at 14-, 21-, and 28-dpi through microCT and histological analysis (Fig 2m). At 14 and 21dpi, timepoints corresponding to the fibrocartilaginous callus development and ossification respectively, the fracture callus of *LysMCre*^*Ert*^*-Camp*^*fl/fl*^ mice exhibited lower BV/TV and apparent density compared with control littermates (Fig 2o-p). In contrast, the callus volume over total volume and callus density were higher in *LysMCre*^*Ert*^*-Camp*^*fl/fl*^ mice at the same timepoints (Fig 2q). These results suggest that the loss of myeloid-derived *Camp* alters the composition and progression of the healing callus.

We next examined fracture healing histologically using Safranin-O/Fast Green stained sections (Fig 2n). At 21dpi, control littermates exhibited a higher proportion of bone within the fracture callus compared with *LysMCre*^*Ert*^*-Camp*^*fl/fl*^ mice, but this difference was no longer apparent at 28 days (Fig 2t). The transient nature of this difference is consistent with a delay in the fracture healing process in the mice globally or conditionally lacking *Camp*. Collectively, these results demonstrate that the presence of *Camp* ensures the proper progression of fracture healing and that myeloid-specific depletion recapitulates the fracture repair phenotype observed following a global knockout.

### Absence of Camp in aged mice results in improved fracture repair

Myeloid-derived *Camp* expression has been studied in a range of disease contexts, including kidney injury and cancer. In the context of bone healing, exogenous CAMP supplementation has been shown to increase osteogenesis *in vitro* and *in vivo*^18, 30, 31^. Despite these observations, the physiologic role of *Camp* in fracture repair remains poorly understood, especially in the context of aging. With our transcriptomic and IHC results, we demonstrate that *Camp* expression is still present throughout the entirety of the healing process in aged mice, just at lower levels compared to their young counterparts. Given that age is an important determinant of fracture repair outcomes, we next wanted to elucidate *Camp*’s function in aged fracture healing as well.

To determine *Camp*’s effect on bone repair, we aged *Camp* global knockout mice and their control siblings 18-to 22-months, corresponding to a human equivalent age of at least 60 to 74 years^32^. After aging the mice, we then performed the tibial fracture and collected at 21dpi, the timepoint where we saw the most significant differences in the young global and conditional depleted mice. Unexpectedly, microCT analysis revealed an increase in BV/TV, apparent density, and callus volume over total volume in fractures from *Camp* knockout mice compared to controls (Fig 3b-d). Safranin-O/Fast Green staining and histomorphometric analysis further demonstrated a significant increase in the proportion of total bone in the fractures from *Camp* deficient mice (Fig 3f). In contrast, aged control mice exhibited a greater proportion of fibrotic tissue compared with *Camp* global knockouts (Fig 3i). Thus, in contrast to the impaired healing observed following *Camp* depletion in young mice, the loss of *Camp* in aged mice was associated with increased bone formation and reduced fibrosis, revealing an age-dependent reversal of the *Camp* fracture healing phenotype.

**Figure 3.**
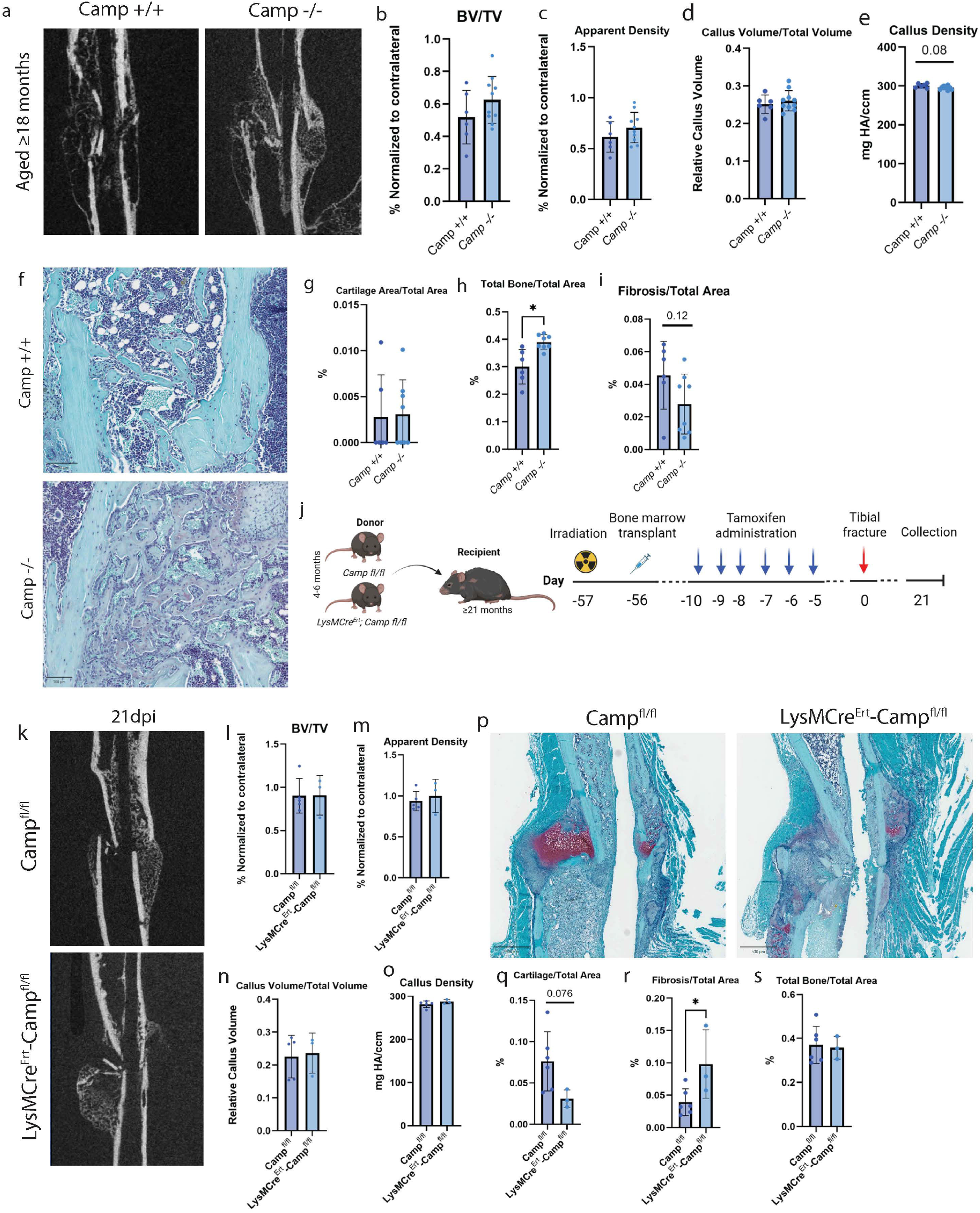
Absence of CAMP is aged mice results in more bone post-fracture. a) Representative microCT images of old (≥ 18 months) *Camp* global knockouts 21 days after a stabilized tibial osteotomy. b-e) Quantification of BV/TV, apparent density, CV/TV, and callus density. f) Representative Safranin-O/Fast Green images. g-i) Quantification of cartilage, total bone, and fibrosis area over total callus area of the old *Camp* global knockout mice. j) Schematic of bone marrow transplants of young (4-6 months) myeloid-specific *Camp* depleted marrow into aged wild type mice. k) Representative microCT images of old control mice post-transplant and fracture. l-o) Quantification of BV/TV, apparent density, relative callus volume (CV/TV), and callus density of recipient mice. p) Representative Safranin-O/Fast Green images. q-s) Quantification of cartilage, total bone, and fibrosis area over total callus area. Results are presented as means ± SD. Statistical significance was determined using 2 tailed Student’s t-test. n ≥ 3, *p < 0.05, **p < 0.01

Young hematopoietic-lineage cells are implicated in the rejuvenation of the pace of fracture repair as demonstrated in heterochronic parabiosis and bone marrow transplants studies^15, 33, 34^. We therefore wanted to examine whether young myeloid-derived *Camp* contributes to the rejuvenation effect of young hematopoietic cells on fracture repair in aged mice. 21-month-old wild type mice underwent total body irradiation (at 500cGy in two separate doses) before being transplanted with marrow from young *Camp*^*fl/fl*^ or *LysMCre*^*Ert*^*-Camp*^*fl/fl*^ donors (Fig 3j). Following transplantation and bone marrow reconstitution, the aged recipient mice underwent the same tamoxifen schedule and tibial fracture procedure as previously described for the young *Camp* conditional depletion experiments. Mice were collected 21-day post-fracture, corresponding to the timepoint at which the most pronounced differences were observed in the young conditional depletion animals.

Histomorphometric analysis revealed that aged mice transplanted with marrow from young *LysMCre*^*Ert*^*-Camp*^*fl/fl*^ exhibited increased proportions of cartilage and fibrosis compared to mice transplanted with control *Camp*^*fl/fl*^ marrow (Fig 3q-s). Thus, the depletion of *Camp* specifically within the young myeloid cells was associated with the persistence of cartilage and fibrotic tissue, suggesting impaired osteogenic progression from the cartilaginous callus to bone. These results would imply that the presence of myeloid-derived *Camp* is necessary for the rejuvenated fracture phenotype as observed in the aged mice that received *floxed* marrow.

### CAMP regulates NLRP3 activation through the P2×7R receptor

To determine which receptor mediates CAMP’s activity during osteogenesis, we evaluated EGFR and P2×7R, known candidate receptors for CAMP, along with their established downstream signaling pathways. Through IHC, we observed no differences in EGFR or P2×7R expression from fracture callus collected from young *Camp* global knockouts or control littermates 7- or 14dpi (Fig 4a; Suppl Fig 4a). Therefore, we then used RT-qPCR analysis to determine whether the expression of the downstream targets was altered in the fracture callus collected from global or conditional *Camp* knockout mice. Of the pathways examined, the fracture calluses collected from global and conditional knockouts showed a consistent decrease of both *Nlrp3* and *Il1b* but not the EGFR targets such as *Jun, Fos*, and *Myc* through the MAPK/ERK or *Nfkb* through the NF-κB pathways (Fig 4b; Suppl Fig 4c-f). Consistent with these findings, Western blot analysis demonstrated reduced NLRP3 protein levels in calluses collected from *Camp* global mutants at 7 days post-fracture (Fig 4c). NLRP3 is a regulator of inflammatory signaling and a known downstream effector of P2×7R^35^. IHC analysis of global Camp mutants and their wild type littermates at 7 days post-fracture demonstrated P2×7R expression in regions of active osteogenesis, with no significant difference in expression between genotypes (Fig 4a). IHC further revealed that NLRP3 expression was concentrated along the perimeter of woven bone at both 7- and 21dpi, overlapping spatially with regions of P2×7R expression (Fig 4e).

**Figure 4.**
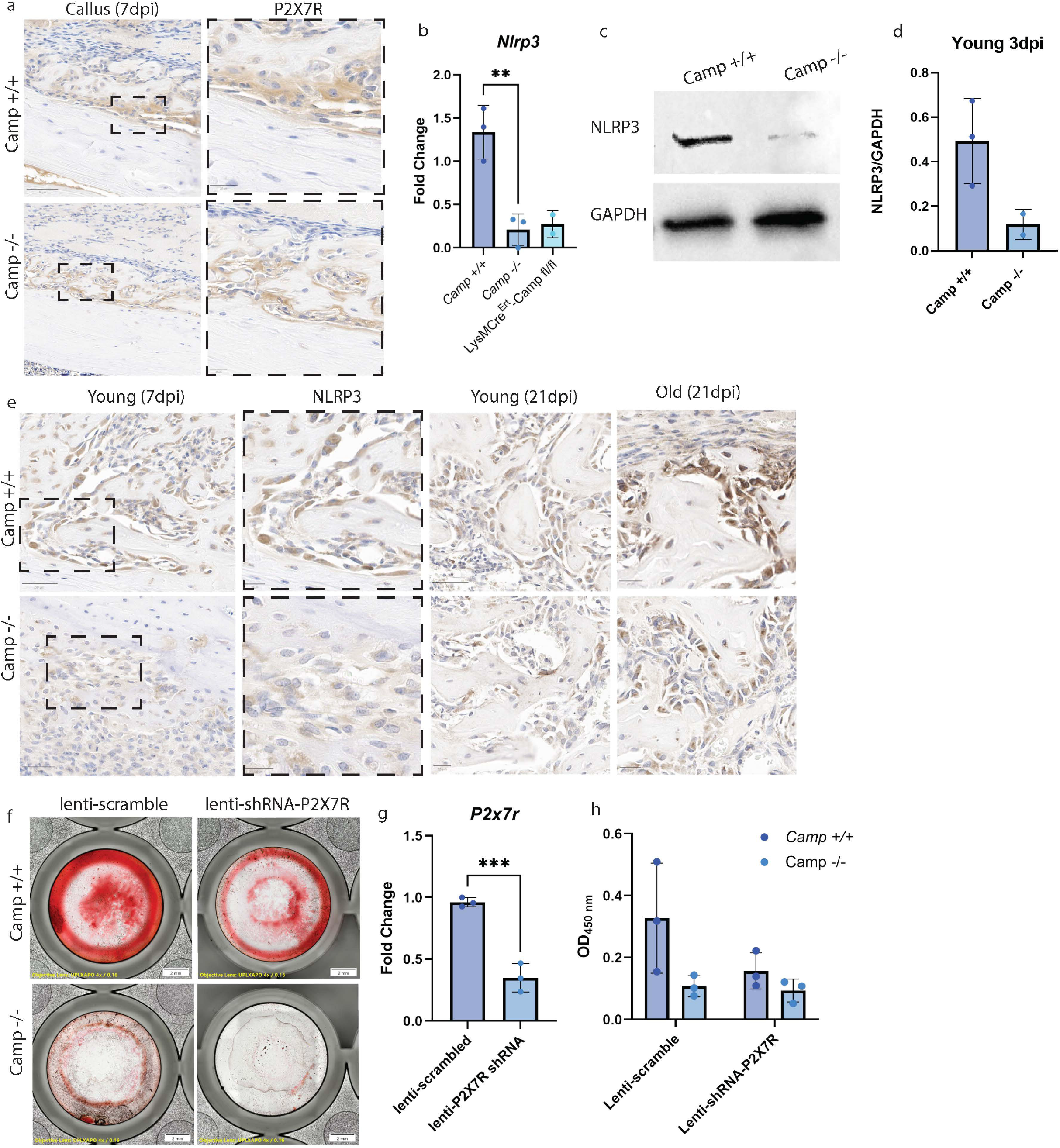
CAMP regulates NLRP3 activation through the P2×7 receptor. a) IHC of P2×7R expression in young *Camp* global knockouts 7 days post-tibial fracture. b) Comparison of *Nlrp3* expression between young *Camp* global, myeloid-specific depletion, and littermate controls 3dpi. c) Representative Western Blot of NLRP3 expression between young *Camp* global knockout and control littermate. d) Quantification of Western Blot. e) IHC of NLRP3 expression from young and old Camp global knockouts 7- and 21dpi. Representative images of BMSCs stained with Alizarin Red after 14 days of osteogenic differentiation after treatment with a scrambled lentivirus or lentiviral shRNA-P2×7R. g) RT-qPCR of *P2×7r* expression after lentiviral transduction. h) Spectrophotometric quantification of Alizarin Red staining. Results are presented as means ± SD. Statistical significance was determined using 2 tailed Student’s t-test. n ≥ 2 *p < 0.05, **p < 0.01, ***p < 0.001

We next investigated whether CAMP activation of P2×7R contributes to downstream osteogenic signaling and whether disruption of either CAMP or P2×7R impairs this response. We used a lentiviral shRNA to knock down *P2×7r* from BMSCs and assessed downstream mineralization by Alizarin Red staining. BMSCs isolated from young wild type mice were cultured with macrophage conditioned media derived from either young *Camp* global mutants or their control littermates (Fig 4f). The BMSCs were then subsequently treated with either a scrambled control lentivirus or a *P2×7r*-targeting shRNA, which reduced *P2×7r* expression by approximately 65% as measured by RT-qPCR (Fig 4g). BMSCs treated with control macrophage conditioned media exhibited robust Alizarin Red staining, whereas exposure to conditioned media from *Camp* deficient macrophages resulted in reduced mineralization (Fig 4h). Notably, the P2×7R knockdown largely eliminated the difference in mineralization between the two conditioned media groups, suggesting that P2×7R signaling is required to the CAMP-dependent osteogenic response. Together, these findings support a model in which macrophage-derived CAMP promotes osteogenic differentiation in BMSCs, at least in part, through P2×7R signaling.

NLRP3 expression is known to be higher in aging in a variety of contexts^36, 37^. While NLRP3 regulates fracture healing in a mouse model of rheumatoid arthritis, its role in the regulation of fracture repair under non-pathological conditions is unknown^38^. To determine the relationship between CAMP and NLRP3 signaling at different ages *in vivo*, we performed IHC on fracture calluses from young and aged global *Camp* knockouts and their respective age-matched control littermates (Fig 4e). As expected, we observed the highest levels of NLRP3 staining in the aged control samples while the aged Camp depleted mice had lower expression level that were comparable to the young controls, raising the possibility that osteogenesis and fracture repair regulation by CAMP is mediated by NLRP3.

In summary, our findings support an age-dependent model of CAMP signaling during fracture repair (Fig 5). In young mice, CAMP promotes osteogenic progression during normal fracture healing. Macrophage-derived CAMP promotes P2×7R-dependent signaling in BMSCs, which is associated with the activation of downstream NLRP3 signaling and osteogenic differentiation. In aged mice, however, the relationship between CAMP signaling and fracture repair is altered. Persistent CAMP expression in the aged fracture microenvironment is correlated with reduced bone formation and increased fibrosis, whereas *Camp* depletion improves these parameters of repair. Moreover, the level of NLRP3, a P2×7R downstream target, correlates with effective osteogenesis and healing, suggesting its role as a mediator in CAMP-regulated fracture repair.

**Figure 5.**
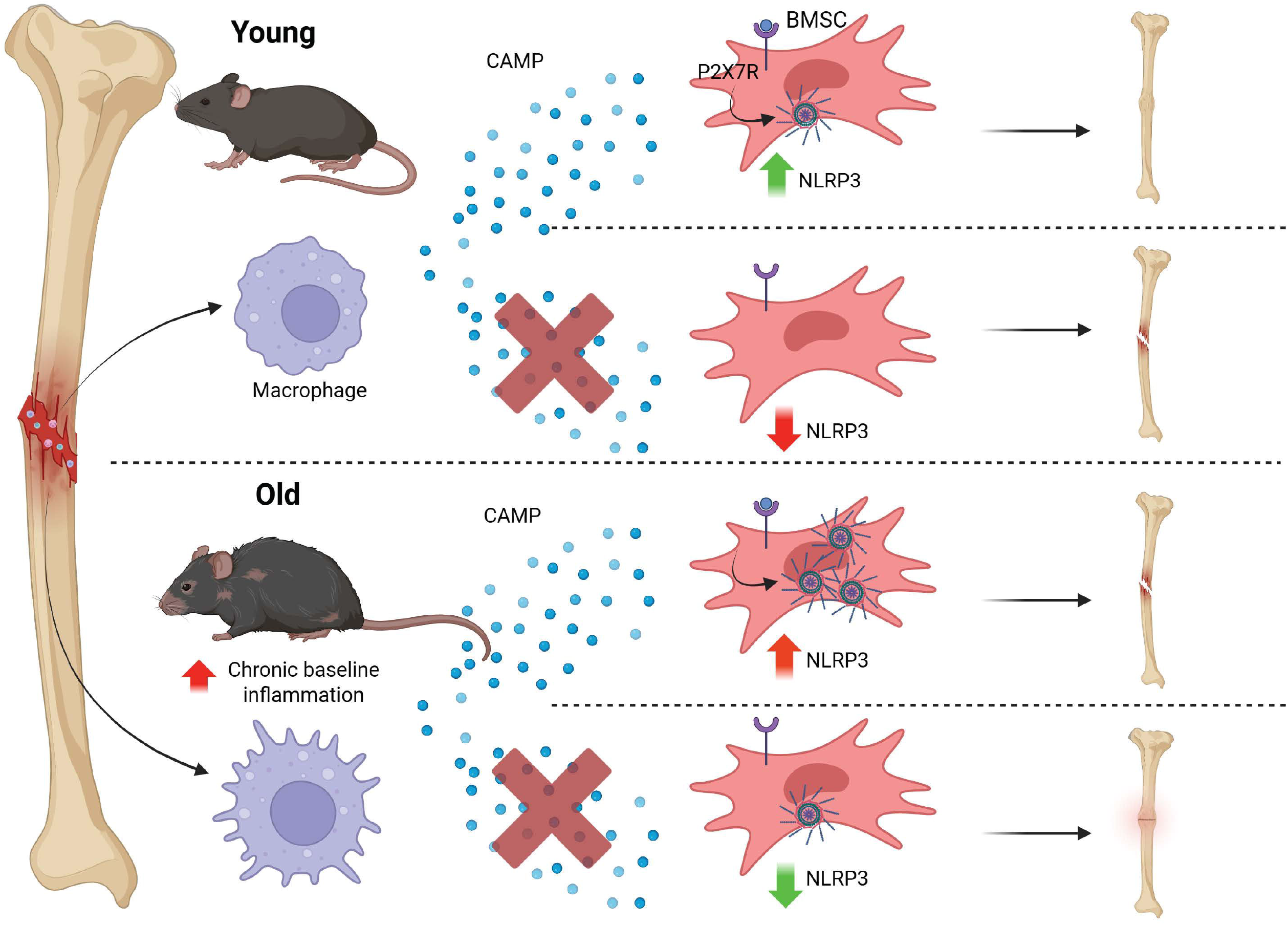
The age-dependent role of CAMP in fracture repair. Schematic demonstrating the different bone fracture outcomes with or without CAMP in both a young and old environment.

## Discussion

Bone fracture repair is a highly coordinated and dynamic process. With age, there is an increased risk of delayed union and nonunion. The early phase of fracture healing is a critical regulator of subsequent tissue repair and represents an opportunity to identify cellular and molecular targets for therapeutic interventions. Aging alters this early inflammatory response, in part through changes in hematopoietic cell composition and a bias toward myeloid differentiation. Using transcriptomic profiling, we identified cathelicidin antimicrobial peptide (*Camp*) as a factor highly expressed by macrophages following fracture in young animals that exerts distinct effects on repair depending on age. Our findings identify *Camp* as an age-dependent regulator of fracture healing and suggest that the functional consequences of inflammatory signaling are strongly influenced by the age-associated healing environment.

*Camp* is a well-established component of the innate immune system, but its role in bone repair has not been extensively characterized. We found that loss of *Camp* delayed fracture repair at earlier healing time points in young mice, whereas *Camp* deficiency in aged mice resulted in calluses with greater bone area. Previous studies have demonstrated that exogenous CAMP supplementation can promote osteogenesis in bone defect models^20, 27^. However, our findings provide evidence that endogenous CAMP has age-dependent effects on fracture repair *in vivo*. These opposing phenotypes further demonstrate how the modulation of inflammatory mediators can have substantially different consequences for tissue repair depending on the biological environment. Our findings also implicate P2×7R and NLRP3 signaling as components of the pathway through which CAMP may influence osteogenic outcomes.

Although CAMP can signal through multiple receptors and pathways, our findings identify P2×7R as the most likely mediator of CAMP-dependent osteogenesis in BMSCs. Targeted P2×7R knockdown *in vitro* impaired the osteogenic response to macrophage-secreted CAMP, supporting a P2×7R-dependent mechanism through which CAMP promotes osteogenesis in BMSCs. Because P2×7R and NLRP3 were not directly manipulated *in vivo*, the fracture healing phenotype observed in our global and myeloid-specific *Camp* depleted mice cannot be definitively attributed to these pathways. Although we did not observe significant and/or consistent change in the downstream targets of the MAPK/ERK and NF-κB pathways, these findings do not exclude additional receptors or mechanisms through which CAMP may regulate fracture repair.

The role for NLRP3 in fracture healing remains incompletely understood. Excessive NLRP3 activation has been associated with impaired fracture healing in a model of rheumatoid arthritis^38^. In a variety of disease contexts, the connection between NLRP3 and inflammation has been well established and has been identified as a therapeutic target to better healing parameters^39-41^. Our data suggests that NLRP3 needs to be at appropriate levels for fracture repair to proceed without a delay; low levels in young *Camp* knockouts or high levels in the aged control mice both resulted in slower healing.

Together, our findings establish CAMP as an age-dependent regulator of fracture repair that further demonstrate the intersection between the early myeloid inflammatory response and downstream osteogenic signaling. The opposing age-dependent effects indicate that CAMP does not have a uniform function in bone healing but instead exerts context-dependent effects that are shaped by the age-associated inflammatory milieu. More broadly, our findings identify the CAMP-P2×7R-NLRP3 axis as a possible mechanistic intermediary between immune signaling and skeletal repair. These results raise the possibility that selective modulation of this pathway could improve fracture healing in older animals while preserving the cellular processes required for bone healing.

## Material and Methods

### Mice

The *Camp* global knockout (*gKO*) mice were purchased from Jackson laboratories. The *LysMCre*^*Ert*^*-Camp*^*fl/fl*^ mice were generated by crossing the *LysMCre*^*Ert*^, also purchased from Jackson laboratories, with the *Camp floxed* mice provided by Dr. Richard Gallo (University of California at San Diego). In all experiments, *Camp gKO* or *LysMCre*^*Ert*^*-Camp*^*fl/fl*^ were compared to their control littermates. All mice were kept in the C57BL/6 background and maintained in a standard, pathogen-free facility of the Division of Laboratory Animal Resources at Duke University according to institutional IACUC protocols. The genotypes of the mice were determined by PCR analysis of genomic DNA extracted from tail clips. *LysMCre*^*Ert*^*-Camp*^*fl/fl*^ depletion was induced by six consecutive doses of tamoxifen at 75mg/kg through intraperitoneal injections then five days of no tamoxifen before the day of fracture surgery.

### Drill Hole Injury

Mice were anesthetized with isoflurane and 5mg/kg buprenorphine. Superficial incision was made to expose the lateral side of the right femur. A 0.8mm diameter bicortical drill hole defect was created using a 21-gauge needle approximately 6mm proximal to the knee joint. The skin was then closed using nylon suture.

### Single Cell RNA preparation, sequencing, and data availability

Femurs from young (2-month) and old (20-month) mice 3 days post-drill hole injury were collected and cleaned of muscle and other extraneous tissue. The bone was washed with 1xPBS before flushing the bone marrow with 1xPBS w/v 0.01% BSA and filtered through a 70um cell strainer. Cell suspensions mixed with reverse transcriptase reagents (Chromium Single Cell Library Kit, 10x Genomics) before loaded into the 10x Genomics Chromium Controller Single-Cell Instrument (10x Genomics). The demultiplexing, barcoded processing, and gene counting and aggregating were completed using Cell Ranger. The data has been uploaded and is available on the NCBI GEO Database with the accession number GSE345479.

### Stabilized tibial fracture

Mice were anesthetized with isoflurane and 5mg/kg buprenorphine. A superficial incision was made to expose the patellar tendon and to insert a 25-gauge needle into the tibial head to make an entry point for the stabilizing insect pin (26001-30, FST). A mid-diaphyseal transverse fracture was made using scissors. Bupivacaine was injected into the open surgical wound before the skin was closed using nylon suture.

### Bone Marrow Transplants

Mice were irradiated at 500cGy (CIX3 Irradiator, Xstrahl) in two separate doses. 24 hours after the irradiation, bone marrow from the donor mice were isolated, depleted of red blood cells, and filtered through a 70um cell strainer with a solution of Hank’s Balanced Salt Solution (HBSS) (14025092, Gibco), fetal bovine serum, and 0.5M EDTA. The resulting cells were then suspended in HBSS at a concentration of 3-6×10^6^ cells in 100uL. Recipient mice were then anesthetized with isoflurane and injected with the cell suspension retro-orbitally. Mice were monitored and given supportive care daily for two months during marrow reconstitution.

### microCT

Limbs were collected at their respective timepoints and placed into 10% neutral buffered formalin. The limbs were fixed for 48 hours at room temperature before being placed in a storage solution of 70% ethanol at 4°C. Samples were taken for microCT scanning and analysis using the vivaCT 80 preclinical system (Scanco, Switzerland). For microCT analysis, X-Ray images were generated and used to identify the fracture line. 1mm above and below the fracture line was analyzed to generate the total volume, bone volume, apparent density, callus volume, and callus density measurements. To account for the changing tissue composition of the healing bone throughout all the timepoints, samples underwent two different thresholding values: a higher one for calcified tissues (termed “bone”) at 480mg HA/cm3 and one for less mature bone (termed “callus”) from 169 to 400mg HA/cm3.

### Immunohistochemistry and histology

Limbs are decalcified in 14% EDTA for 14 days. The samples were then paraffin embedded and sectioned at 5-micron thickness. After baking overnight at 60°C, slides were taken for Safranin-O/Fast Green staining to quantify cartilage and bone area or immunochemistry (IHC) for CAMP (12009-1-AP, 1:250, Proteintech) and NLRP3 (PA5-79740, 1:1000, Invitrogen). For IHC, the primary antibodies were incubated overnight at 4°C before development with rabbit IHC kit (PK-6101,Vector Labs). Nuclei were stained with hematoxylin. For quantification of positive immunohistochemical signal, slides were scanned and analyzed using an automated QuPath software for cell detection using four different 10,000 um^2^ regions of interest within the callus for technical replicates per biological sample.

### RT-qPCR analysis

Fractured limbs were dissected and cleaned of soft tissue. Calluses were isolated by trimming off the tibial head and ankle before being flash frozen in liquid nitrogen. Using a stainless-steel pestle and mortar that was pre-chilled with liquid nitrogen, the calluses were then manually homogenized into powder. Total RNA was collected through TRIzol (Invitrogen) and an isolation kit (R2052, Zymo) according to the kit protocol. cDNA synthesis was performed using 1ug of total RNA and iScript (1708841, Biorad). RT-qPCR quantification was performed using SYBRGreen (4364344, Applied Biosystems) on a QuantStudio 6 Flex System (Thermo) and the relative mRNA expression levels were normalized to the ACTB expression levels using the Δ ΔCT method. Camp-forward primer; CTGTGGCGGTCACTATCACT, Camp-reverse primer; GTTCCTTGAAGGCACATTGC, Nlrp3-forward primer; TCACAACTCGCCCAAGGAGGAA, Nlrp3-reverse primer; AAGAGACCACGGCAGAAGCTAG, Nfkb-forward primer; GCTGCCAAAGAAGGACACGACA, Nfkb-reverse primer; GGCAGGCTATTGCTCATCACAG, Il1b-forward primer; TGGACCTTCCAGGATGAGGACA, Il1b-reverse primer; GTTCATCTCGGAGCCTGTAGTG, cMyc-forward primer; TCGCTGCTGTCCTCCGAGTCC, cMyc-reverse primer; GGTTTGCCTCTTCTCCACAGAC, Fos-forward primer; GGGAATGGTGAAGACCGTGTCA, Fos-reverse primer; GCAGCCATCTTATTCCGTTCCC, Jun-forward primer; CAGTCCAGCAATGGGCACATCA, Jun-reverse primer; GGAAGCGTGTTCTGGCTATGCA, ActB-forward primer; GGCTGTATTCCCCTCCATCG, ActB-reverse primer; CCAGTTGGTAACAATGCCATGT.

### Western blotting analysis

Fractured limbs were dissected and cleaned of soft tissue. Calluses were isolated by trimming off the tibial head and ankle before being flash frozen in liquid nitrogen. Using a pre-chilled stainless-steel pestle and mortar, the calluses were then manually homogenized into powder. Protein lysates were then made by adding RIPA buffer (89900, Thermo) with anti-phosphatase (A32957, Thermo) and anti-protease (11836170001, Roche) to the powdered calluses. Samples were then agitated and centrifuged before collecting the resulting supernatant. Cell lysates were separated by SDS-PAGE on precast 4-15% polyacrylamide gels (4568084, Biorad) and blotted onto 0.45um PVDF membranes (88518, Thermo). The membranes were incubated overnight at 4C with antibodies against CAMP (12009-1-AP, 1:500, Proteintech), NLRP3 (PA5-79740, 1:500, Invitrogen), GAPDH (2118S, 1:1000, Cell Signaling), and Vinculin (MAB3574, 1:1000, Sigma). The membranes were then re-probed with the appropriate horseradish peroxidase secondary antibodies at 1:1000 concentration. The blots were visualized via ECL (1705061, Biorad) or super ECL (PI34577, Thermo) and ChemiDoc imaging system (Biorad) before being quantified with ImageJ.

### Macrophage conditioned media

Hindlimbs from *Camp WT* or *gKO* were dissected and the soft tissue was removed. The heads of the tibia and femur were trimmed off and placed into a 0.5mL Eppendorf tube that was punctured with an 18-gauge needle. The 0.5mL tube with the limbs is then placed in a 1.5mL tube and centrifuged for 10 seconds at 10,000xg. The bone marrow underwent red blood cell lysis for 4 minutes and then strained through a 70um strainer. Cells were then stained with Live/Dead NIR, CD45 FITC, and F4/80 APC for later sorting for only Live CD45+ F4/80+ macrophages. After sorting, the cells were washed 3 times with sterile 1xPBS before being plated at 1×10^6^ on a 15cm plate with aMEM media with 20% FBS and 1% penicillin/streptomycin. After another 24 hours, the media was collected. The media was syringe filtered with 0.22um filter and frozen in 5mL aliquots at -20°C until use.

### Osteogenic differentiation assay

Hindlimbs from wild type mice were dissected and the soft tissue was removed. The heads of the tibia and femur were trimmed off and placed into a 0.5mL Eppendorf tube that was punctured with an 18-gauge needle. The 0.5mL tube with the limbs is then placed in a 1.5mL tube and centrifuged for 10 seconds at 10,000xg. The bone marrow underwent red blood cell lysis for 4 minutes and then strained through a 70um strainer. Cells were then counted and plated at a density of 1.74×10^5^ in a 48 well plate. The cells were kept in hypoxic conditions for 7 days for cell expansion with aMEM media with 20% FBS and 1% penicillin/streptomycin. Then moved to a 1:1 mixture of macrophage conditioned media and osteogenic differentiation media (aMEM with 10% FBS, 1% penicillin/streptomycin, 8mM b-glycerol phosphate, 50uM ascorbic acid, and 0.1uM dexamethasone) and normoxic conditions for 14 days. Media was changed every 2-3 days during this period.

### Lentivirus transfection and transduction

Lentiviral vectors, scramble-shRNA control (VB010000-9526zpu) and shRNA for mouse *P2rx7* (VB900222-1117ryb), were purchased from VectorBuilder. The plasmids were grown in LB broth with 10mg/mL of ampicillin overnight at 37C. The DNA was purified and collected using a midiprep kit (D4201, Zymo). 298T cells were plated and allowed to grow to 40-60% confluency. Transfection of the cells was conducted using 3ug of transgene, 2ug of pCMV-dR8.2 (8455, Addgene), and 1ug of pCMV-VSV-G (8454, AddGene) with optiMEM (31985062, Gibco) and GeneJuice (70967, Sigma). After 48 hours, the media was collected and syringe sterilized with a 0.45um filter. Transduction was done with 10ug/mL polybrene in the final 24 hours of hypoxic expansion of the bone mesenchymal stromal cells with basal culture media. When cells were moved to osteogenic and macrophage conditioned media and normoxic conditions, 2mg/mL puromycin was added during each media change to ensure cells without lentiviral transduction were removed.

### Alizarin Red staining and quantification

Alizarin red solution was made at 2% with distilled water and pH adjusted to 4.1-4.3. Plates were rinsed with sterile 1xPBS prior to staining. Wells were then incubated with 10% formalin for 15 minutes, rinsed with distilled water, incubated with the staining solution for 15 minutes at room temperature, rinsed with water again, and then dried. Once the plates were fully dried, wells were then imaged (APX100, Evident). For quantification, 800uL of 10% acetic acid was added to each well and incubated at room temperature on a shaker for 30 minutes. After incubation, a cell scraper was used to further homogenize the monolayer before transferring the solution to an Eppendorf tube. The solution was then vortexed for 30 seconds before being incubated at 85°C for 5 minutes. After cooling to room temperature, the solution was centrifuged at 20,000g for 15 minutes. 500uL of the supernatant is transferred to a new Eppendorf and neutralized with 200uL of 10% ammonium hydroxide solution. 150uL of the supernatant is then plated in triplicate in a 96 well plate and read at 405nm (SpectraMax i3x, Molecular Devices).

### Statistical analysis

All data are represented as the mean ± standard deviation. Statistical differentiates were analyzed using two-tailed Student’s t-test or Welch’s t-test where appropriate. Differences of p ≤ 0.05 were considered statistically significant. Minimum sample size was determined using power analysis from previous publications.

## Acknowledgements

We thank Duke University’s Division of Laboratory Animal Resources staff for the care of our animals and DCI Flow Cytometry core staff Dr. Bin Li, Lynn Martinek, and Yingqiu Guo for their technical assistance. We thank Dr. Richard Gallo of University of California-San Diego for providing the *Camp floxed* mice.

This study was supported by the National Institute on Aging (R01AG072058) and the National Institute of Child Health and Human Development (T32HD040372).

## Author Contributions

Nguyen, T., Yahara, Y., and Alman, B. conceived and planned the experiments for the study. Nguyen, T. carried out the experiments, analyzed the results, and wrote the manuscript. Shimada, E. assisted in the data processing and troubleshooting of the single cell RNA sequencing. Puviindran, V. provided feedback on experimental design and guidance on immunohistochemistry, RT-qPCR, Western Blot, and lentiviral transfection. Nadesan, P. supported mouse maintenance, surgery, and dissection. Ishikawa, K. and Marius, C. helped with data analysis and interpretation. Alman, B. supervised the project.

