## Supplemental Figures for "Age-Dependent Effects of Macrophage Derived Cathelicidin Antimicrobial Peptide on Bone Fracture Repair"

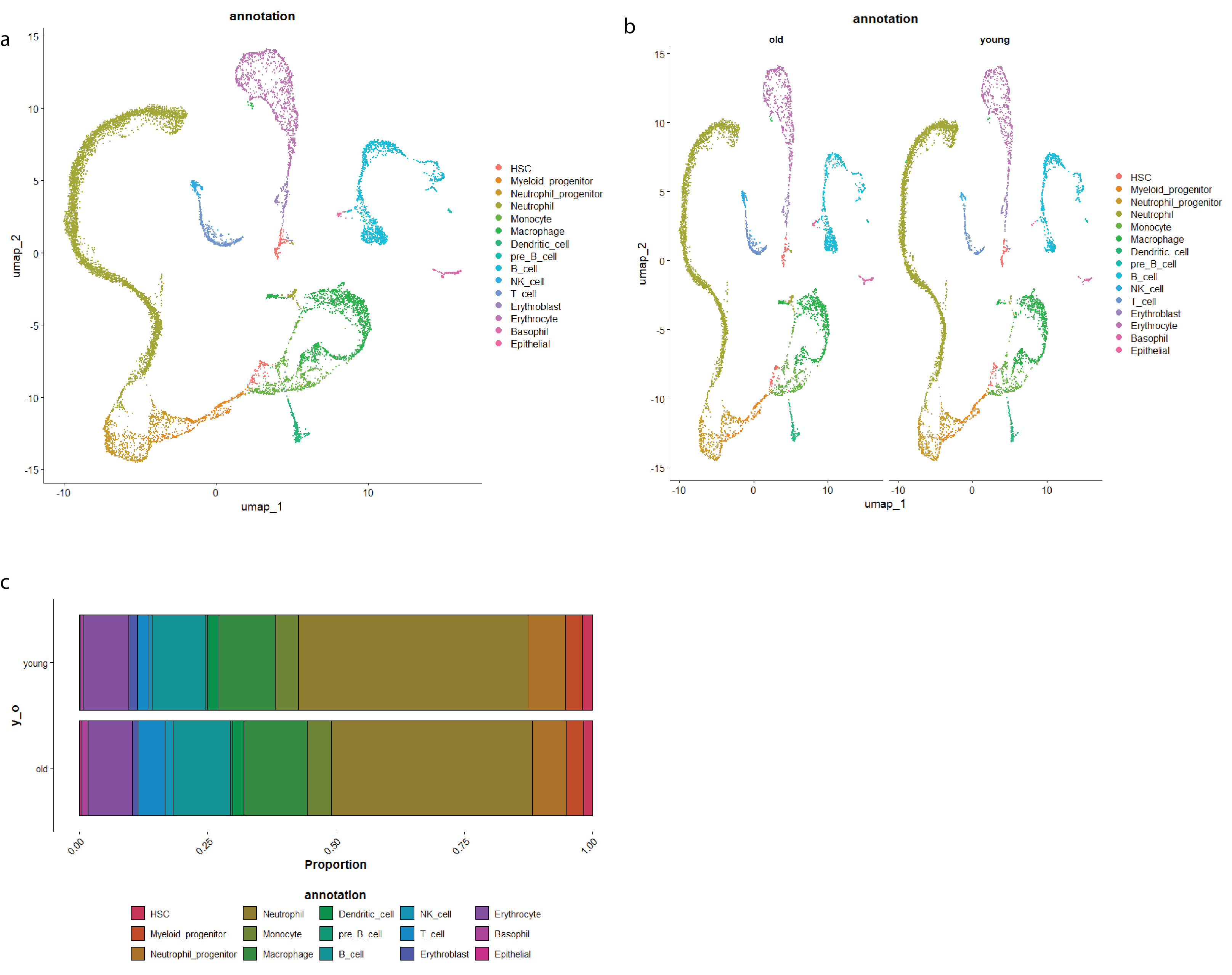

Supplemental Figure 1. Hematopoietic lineage cells have age related differences 3 days following drill hole injury. a) UMAP of integrated dataset of young mice (2 months) and old mice (20 months) collected 3 days after a bicortical femoral drill hole injury b) Comparison of age-specific cell population contribution to the integrated UMAP. c) Comparison of cell type proportion according to age.

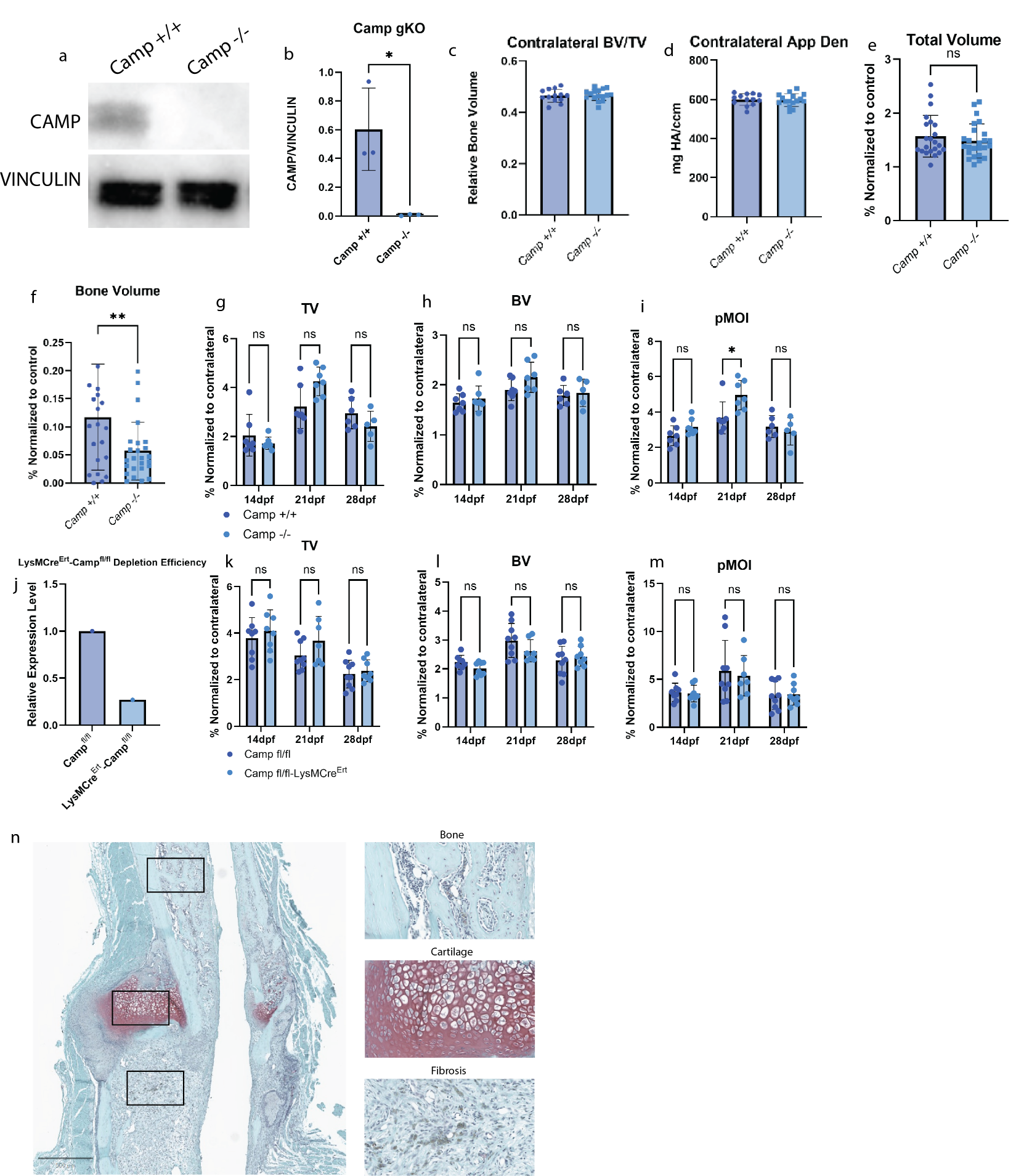

Supplemental figure 2. Validation of young *Camp* global and conditional knockout mice and histomorphometric parameters a) Western Blot validation of *Camp* global knockout mice of splenic protein lysate. b) Quantification of Western Blot. c-d) Relative bone ratio and apparent density of the contralateral limb of *Camp* global knockout mice and their control littermates from drill hole injury experiment. e-f) Total volume and bone volume of the injured drill hole limb normalized to the contralateral of *Camp* global knockout. g-i) Total volume, bone volume, and pMOI of *Camp* global knockout mice and their control littermates after tibial fracture j) qPCR of *Camp* from CD45+F4/80+ cells pooled from 3 mice after tamoxifen administration. k-m) Total volume, bone volume, and pMOI of *LysMCre^Ert^-Camp^fl/fl^* conditional knockout mice and their control littermates after tibial fracture. n) Representative images of bone, cartilage, and fibrosis as visually defined for histomorphometry. Results are presented as means ± SD. Statistical significance was determined using 2 tailed Student’s t-test. *p < 0.05, **p < 0.01

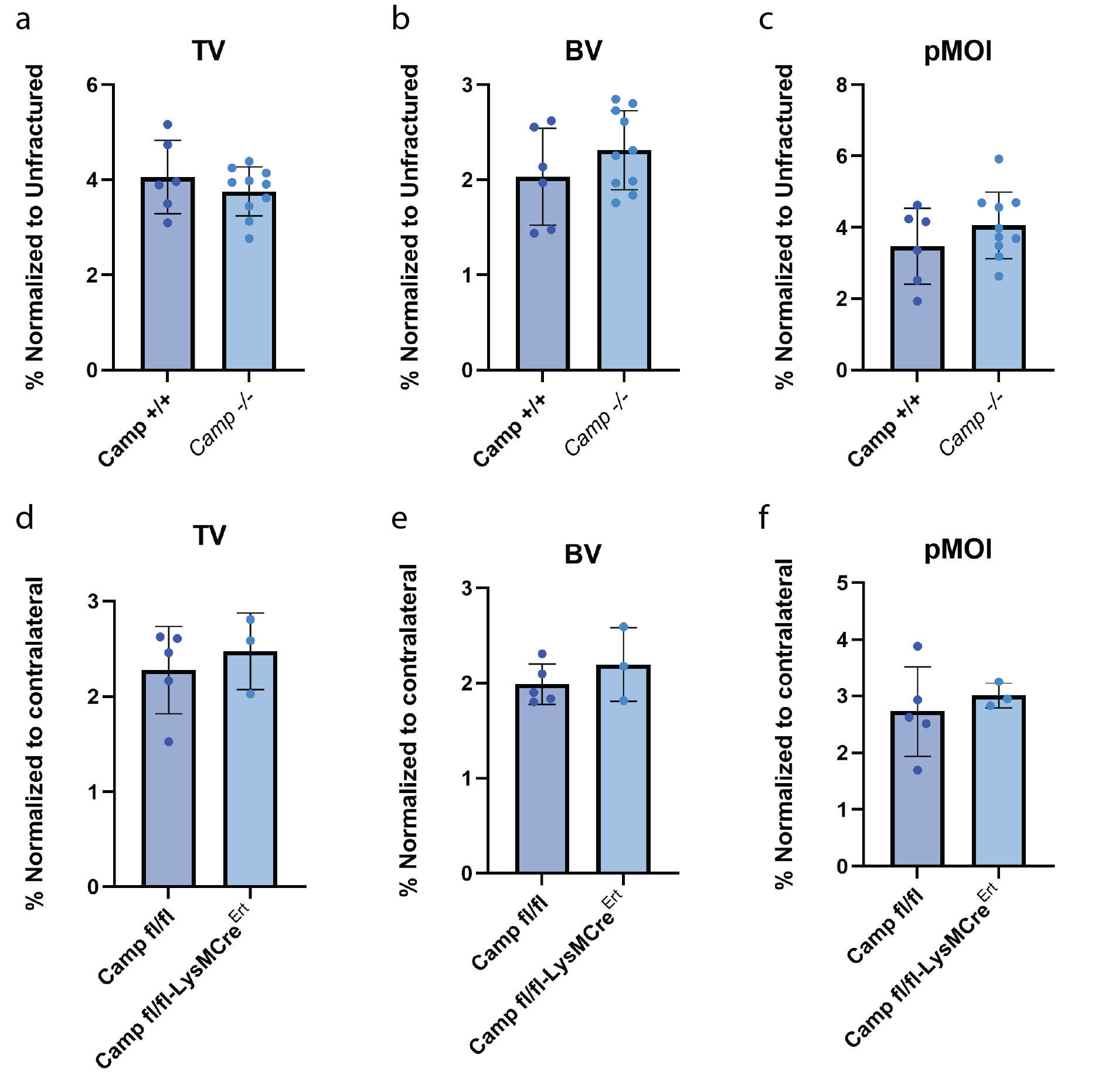

Supplemental Figure 3. Other microCT parameters of aged and bone marrow transplant experiments demonstrate no significant changes. a-c) Total volume, bone volume, and pMOI of old (≥ 18 months) *Camp* global knockouts 21 days after tibial fracture. d-f) Total volume, bone volume, and pMOI of aged bone marrow recipients 21 days after tibial fracture. Results are presented as means ± SD. Statistical significance was determined using 2 tailed student’s t-test. n ≥ 3

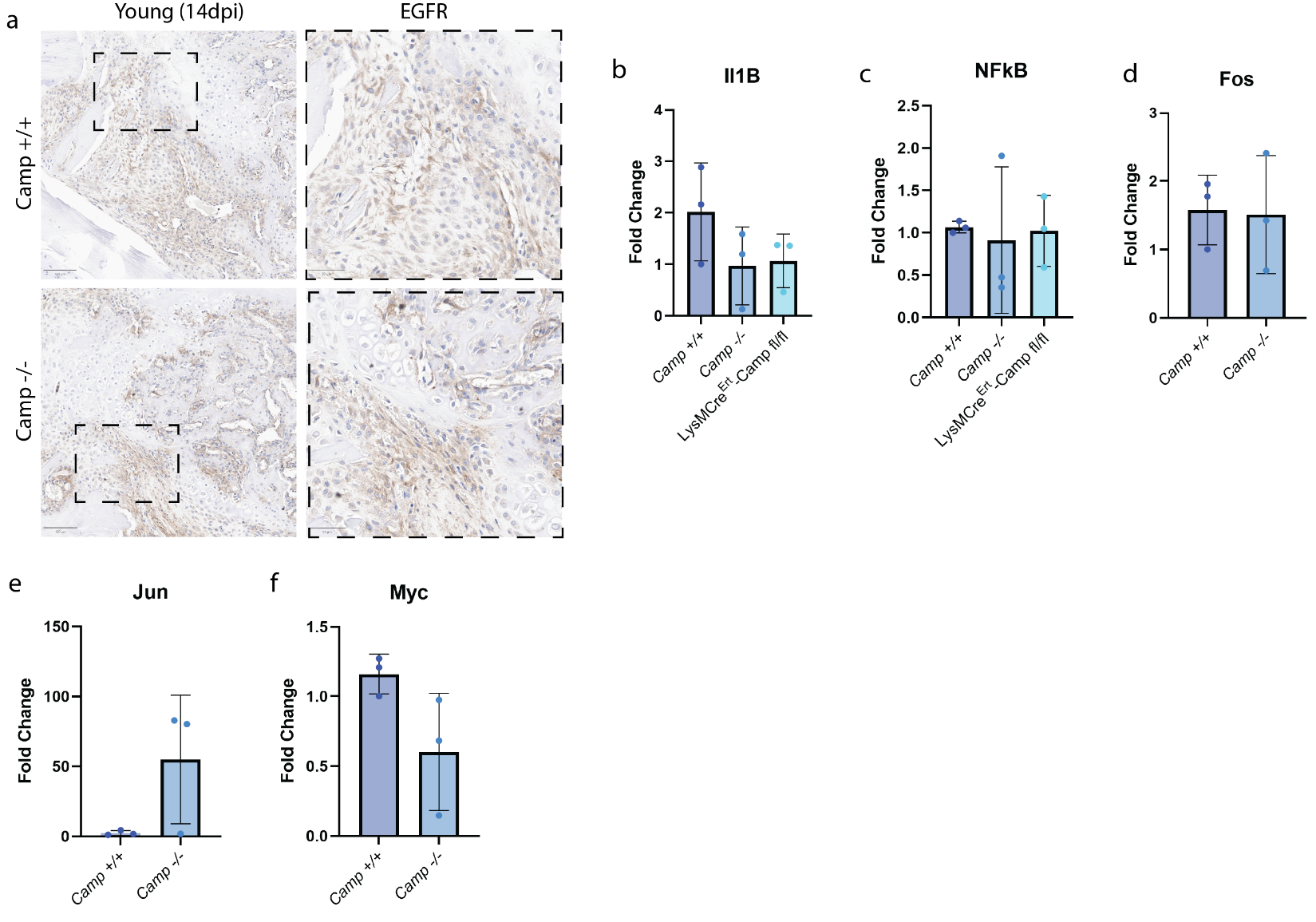

Supplemental Figure 4. EGFR and downstream candidate pathways, MAPK/ERK and NFκB, demonstrated no significant changes. a) IHC of EGFR expression in young *Camp* global knockouts 14 days post-tibial fracture. b-c) qPCR results of *Il1b* and *Nfkb* of young *Camp* global, myeloid-specific depletion, and littermate controls 3 days post-tibial fracture. d-f) qPCR results of *Fos*, *Jun*, and *cMyc* of young *Camp* global knockouts and control littermates 3 days post-fracture. Results are presented as means ± SD. Statistical significance was determined using 2 tailed student’s t-test. n ≥ 3
